# The moonlighting functions of SEC13 in nonsense-mediated mRNA decay and the regulation of ER stress

**DOI:** 10.64898/2026.05.01.722270

**Authors:** Laura Monaghan, Chetan Srinath, Abigail R. Mann, Graeme R. Grimes, Dasa Longman, Javier F. Cáceres

**Affiliations:** MRC Human Genetics Unit, Institute of Genetics and Cancer, University of Edinburgh, Edinburgh EH4 2XU, United Kingdom

**Keywords:** RNA-quality control, nonsense-mediated decay (NMD), SEC13, UPR, stress response, moonlighting

## Abstract

The nonsense-mediated decay pathway (NMD) is an RNA quality control mechanism that regulates the stability of target RNAs. We previously identified the ER-localized SEC13 protein as a novel NMD factor in *C. elegans* and in HeLa cells; raising the possibility that it could be involved in regulating the stability of mRNAs translated at the ER. SEC13 is a component of several cellular complexes, including the COPII vesicle coat, the nuclear pore complex (NPC) and the nutrient sensing GATOR2 complex. Here, we show that SEC13 interacts with core NMD factors and using a newly developed dual-color fluorescent NMD sensor in U2OS cells, we assessed SEC13 NMD activity, at a single-cell level. Transcriptomic profiling revealed that unlike the previously described ER-NMD factor, NBAS, SEC13 co-regulates the stability of substrates translated both in the cytoplasm and at the ER. We also show that SEC13 function in NMD is largely independent of its function in other cellular complexes. Altogether, these results show that SEC13 is a bona fide NMD factor in mammalian cells. Finally, we utilized an ER stress-activated indicator (ERAI) in U2OS cells to demonstrate that SEC13, together with canonical NMD factors, has a role in the regulation of the unfolded protein response (UPR) at the ER. Thus, the moonlighting functions of SEC13 include a role in NMD pathway and the regulation of ER stress.

## INTRODUCTION

Nonsense-mediated decay (NMD) is an evolutionary conserved RNA quality control mechanism with a central role in gene expression. NMD targets mutated mRNAs harboring premature termination codons (PTCs) for degradation, preventing the accumulation of truncated proteins that could be toxic for the cell (Kurosaki et al. 2019; Monaghan et al. 2023). This has an effect in the modulation of the phenotypic outcome of human diseases caused by frameshift or nonsense mutations that introduce PTCs (Bhuvanagiri et al. 2010; McMahon and Maquat 2025). The NMD pathway also controls the stability of non-mutated cellular transcripts and regulates different physiological processes, including neural development and the stress response (Goetz and Wilkinson 2017; Tan and Wilkinson 2025; Karousis and Mühlemann 2022).

Mechanistically, NMD is intimately linked to mRNA translation and is triggered by the recognition of a PTC by the elongating ribosome. A key factor in this RNA surveillance mechanism is the ATP-dependent RNA helicase of the SF1 superfamily UPF1 (Kim and Maquat 2019). First, a surveillance complex, termed SURF, comprising the ribosome release factors, eRF1 and eR3, UPF1, and its associated kinase, SMG1 is assembled (Kashima et al. 2006). Secondly, interactions of this surveillance complex with core NMD factors, UPF2 and UPF3, and with an exon junction complex (EJC) leads to UPF1 phosphorylation and the assembly of a decay-inducing complex (DECID) followed by the recruitment of mRNA degrading activities (Embree et al. 2022; Le Hir et al. 2001; Boehm et al. 2021).

We previously carried out RNAi screens in *C. elegans*, which identified two novel NMD factors localizing to the endoplasmic reticulum (ER) that are also essential for viability, suggesting that they fulfil additional non-NMD functions in nematodes, where the NMD pathway is not essential. These include *smgl-1* corresponding to the human gene *NBAS* (neuroblastoma amplified sequence), and *npp-20*, the human homolog *SEC13* (Longman et al. 2007; Casadio et al. 2015). We went on to characterize NBAS as a bona fide NMD factor that acts together with UPF1 to regulate NMD in nematodes, zebrafish and human cells (Anastasaki et al. 2011; Longman et al. 2013). More recently, we showed that NBAS recruits UPF1 to the membrane of the ER and activates a local NMD response, which we termed ER-NMD, targeting for degradation mRNAs translated at the ER (Longman et al. 2020).

SEC13 is a short WD40 domain-containing protein that is a component of several cellular complexes. It forms a heterotetramer with SEC31 to assemble the outer coat of the COPII (Coat Protein Complex II) vesicle complex with a role in ER-to-Golgi anterograde transport (Barlowe et al. 1994; Stagg et al. 2006; Lord et al. 2013). It is also part of a large sub-complex with nine additional proteins forming the outer ring of the nuclear pore (Siniossoglou et al. 1996), and forms a complex with SEH1L to structurally stabilize the nutrient sensing GATOR2 complex, which is a regulator of mRNA translation through mTORC1 (Valenstein et al. 2022). Accordingly, SEC13 has been shown to localize mainly to the ER and to the nuclear pore complex (NPC), and to function as a nucleo-cytoplasmic shuttling protein (Enninga et al. 2003). Thus, SEC13 is an example of a moonlighting protein, with a variety of unrelated cellular functions (Copley 2014).

We previously uncovered some evidence that shows that SEC13 is involved in the NMD pathway in human cells (Casadio et al. 2015). Its localization to the ER suggested that, like NBAS, SEC13 could potentially be involved in the ER-NMD pathway. Here, we aim to elucidate the mechanistic roles of SEC13 in NMD and to understand the multi-faceted functions of this versatile protein in mammalian cells. Importantly, we show a role for SEC13 in the regulation of the NMD response of mRNAs translated both at the ER and in the cytoplasm. We present evidence that highlights a role for SEC13 in the NMD response, which operates independently of its role as a member of other cellular complexes. We also demonstrated a role for SEC13 and other canonical and ER-NMD factors in the regulation of the stress response at the ER.

## RESULTS

### The multifunctional SEC13 protein interacts with core NMD factors

SEC13 is a known component of three different cellular complexes with diverse subcellular localization and unique roles. To characterize the involvement of SEC13 in the NMD pathway, we searched for SEC13 interactors. We used CRISPR/Cas9 genome editing in HeLa cells to introduce an eGFP/3xFlag-tag at the C-terminus of the endogenous SEC13 locus (HeLa-SEC13-tag) (Fig. 1A), selecting this cell type as we previously carried out interactome studies for ER-NMD factor, NBAS, in this cell line (Longman et al. 2020). We carried out immunoprecipitation (IP) and mass spectrometry (MS) of anti-GFP SEC13, which resulted in the identification of multiple components of the nuclear pore complex (NPC), COPII vesicles and the GATOR2 complex, all predicted interactors of SEC13 (Fig. 1A; Supplemental Table S1). For instance, amongst the COPII interactors, we detected SEC16A, which agrees with previously published data, reporting an interaction between the two proteins (Hughes et al. 2009; Whittle and Schwartz 2010). We did not observe an enrichment of NMD factors, which could suggest that, if these interactions occur, they are of a more transient nature, as we previously have shown with the ER-NMD factor, NBAS in HeLa cells (Longman et al. 2020).

**FIGURE 1.**
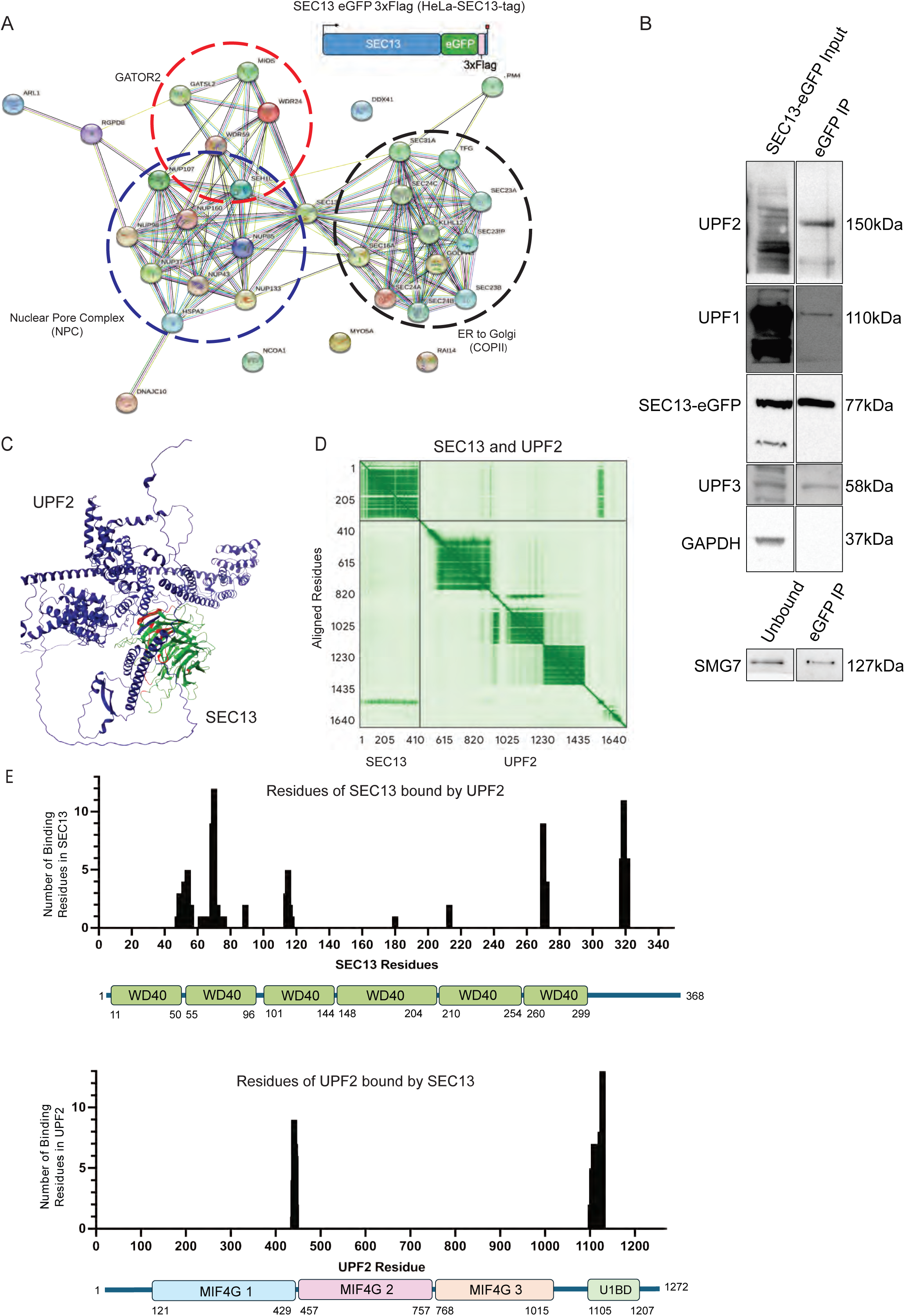
SEC13 is part of different cellular complexes and interacts with core NMD factors*. (A)* Cartoon depicting the endogenously tagged SEC13 locus in HeLa cells, including a C-terminal 3xFLAG/eGFP tag (HeLa-SEC13-Tag, upper panel). String network of interacting proteins identified by mass spectrometry of anti-GFP Immunoprecipitation (lower panel). SEC13 interacts with protein complexes involved in the GATOR2 complex, the nuclear pore complex (NPC) and the COPII complex. A full list of identified interacting proteins is presented in Supplemental table 1. *(B)* Endogenously tagged SEC13 was immunopurified with anti-GFP antibody and the co-immunoprecipitation of core NMD factors was analyzed by Western Blot with corresponding antibodies. *(C)* AlphaFold3 analysis of SEC13 with UPF2 suggests a potential direct interaction (SEC13 in green; UPF2 in blue; interacting residues highlighted in red). *(D)* Predicted Aligned Error (PAE) plot identified internal structure with highlighted area of hypothesized interactions between UPF2 and SEC13. *(E)* Full length SEC13 displaying the six WD40 domains and showing the potential interacting residues with UPF2 (upper panel) and vice versa (lower panel).

To probe further for the interactions of SEC13 with canonical NMD factors, we followed a candidate approach, where we immunoprecipitated SEC13 with GFP-coupled magnetic beads, and probed for the presence of NMD interactors by Western blot analysis. We observed that endogenous SEC13 interacts with several canonical NMD factors, including UPF1, UPF2 and UPF3 (Fig. 1B). SEC13 was also previously shown to interact with the core NMD factor SMG7, which forms a complex with SMG5 leading to RNA degradation (Loh et al. 2013). We confirmed the interaction of SEC13 with SMG7, suggesting that SEC13 remains associated with NMD complex at later stages (Fig. 1B, lower panel).

To model the direct interaction of SEC13 with canonical NMD factors, we carried out AlphaFold structural modelling, using AlphaFold3 (AF3) (Abramson et al. 2024). As a positive control, we included the COPII factors, SEC31A/B, which together with SEC13 form a heterotetramer (Stagg et al. 2006). As expected, a direct interaction of SEC13 and SEC31 was predicted in all five models with binding across the WD40 domains of SEC13 (Supplemental Fig. S1A, B). Focusing on canonical core NMD factors, this analysis suggested a direct interaction of SEC13 with UPF2 in 3 of the 5 models (Fig. 1C, D). The major areas of binding in SEC13 centered on the second WD40 domain with a small amount of binding at the C-terminus, which are predicted to form a binding pocket in the 3D structure. The predicted interaction of SEC13 with UPF2 is primarily localized in the early residues of the UPF1 binding domain at the C terminus of UPF2, which again is predicted to form a binding pocket with the small region between the MIF4G 1 and 2 (Fig. 1E). This highlights a potential for SEC13 to influence the binding of UPF2 with UPF1, and the assembly of the NMD complexes. By contrast, this approach did not reveal any predicted direct interactions of SEC13 with either UPF1 (Supplemental Fig. S1C), UPF3 or other NMD factors (data not shown).

We next used UPF1 phosphorylation as a diagnostic tool to infer the timing of SEC13 recruitment to the NMD complex. The C126S UPF1 mutant resembles the hypophosphorylated state of UPF1 that is present in the early surveillance (SURF) complex, whereas the K498A UPF1 mutant mimics the hyperphosphorylated UPF1 present in the late decay-inducing (DECID) complex (Kashima et al. 2006; Weng et al. 1996). SEC13 bound both UPF1 mutants, with enrichment for C126S UPF1, suggesting SEC13 is recruited during the early stages of NMD, and remains associated following the transition to the DECID complex (Supplemental Fig. S1D). The binding of SEC13 to the hypophosphorylated UPF1 mutant is consistent with previous observations for the ER-NMD factor NBAS (Longman et al. 2020), and the RNA helicase DHX34 (Hug and Cáceres 2014), perhaps suggesting that most of the regulatory steps occur in the early stages of the NMD response. Altogether, these experiments confirm that SEC13 is part of different cellular machineries, including the GATOR2, the NPC and COPII complexes, and interacts with core NMD factors UPF1, UPF2, UPF3 and SMG7. This together with AlphaFold structural modeling suggests that SEC13 is bound directly to UPF2, in a larger complex with UPF1 and UPF3.

### SEC13 localizes to the endoplasmic reticulum (ER) and is in the proximity of NMD factors

To investigate SEC13 subcellular localization, we introduced an FKBP12^F36V^/3xFLAG/eGFP-tag at the C-terminus of the endogenous *SEC13* locus in U2OS osteosarcoma cells (U2OS-SEC13-tag), using CRISPR/Cas9-mediated genome editing (Fig. 2A, upper panel). We switched to U2OS cells, to align with the NMD sensors integrated in this cell line (see below). Using this cell line harboring an endogenously tagged *SEC13* locus, allowed us to assess the subcellular localization of endogenous SEC13 via eGFP (Figs. 2B, C), as well as to use the FLAG tag for immunoprecipitation studies, and the FKBP12^F36V^ to induce acute degradation the protein (Supplemental Fig. S2A, B).

**FIGURE 2.**
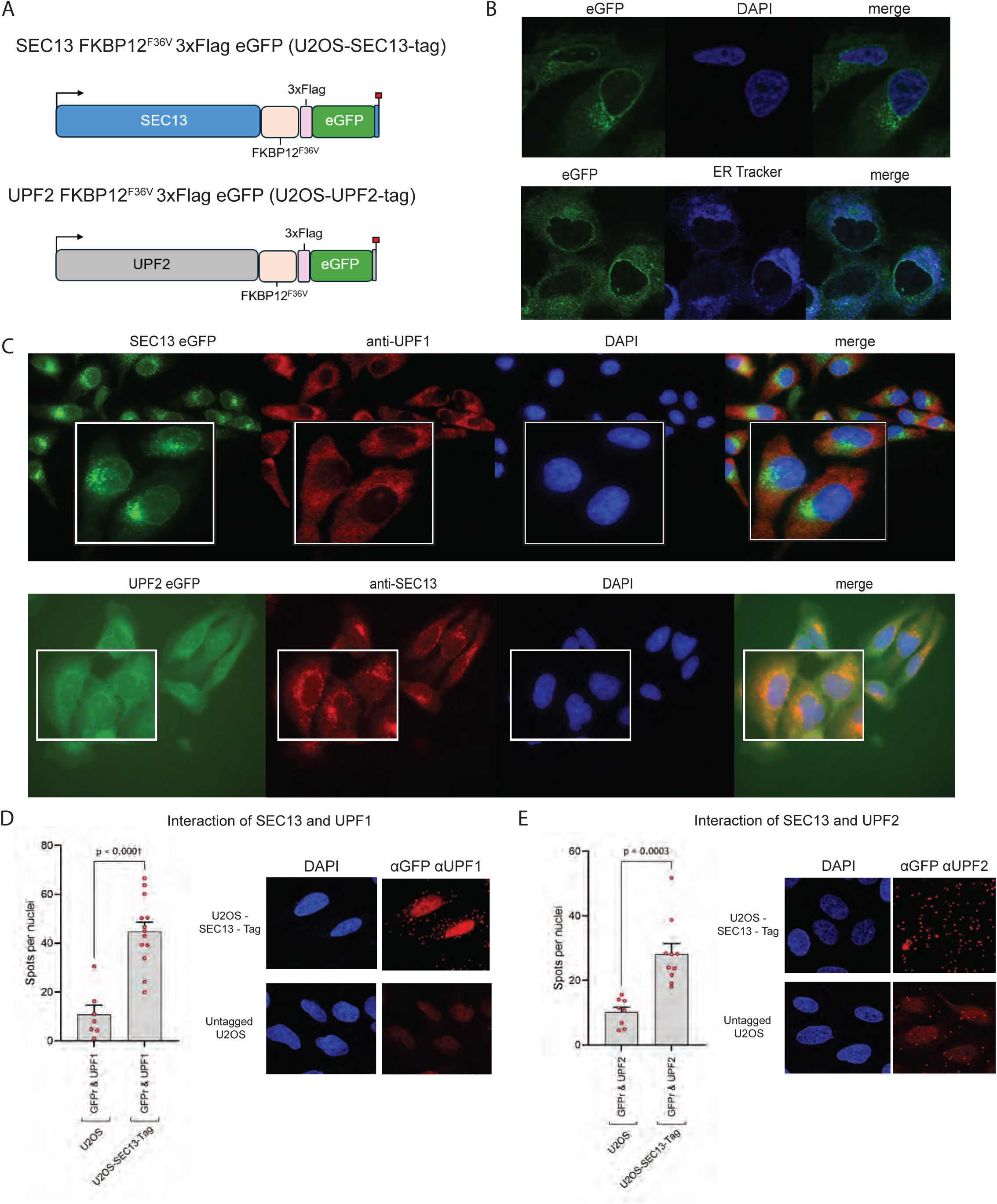
SEC13 localizes to the ER in proximity to NMD factors. *(A)* A C- terminal tag, comprising an FKBP12^F36V^ sequence followed by 3xFLAG/eGFP was inserted at the C-terminus of the *SEC13* (upper panel) and *UPF2* loci (lower panel), in U2OS osteosarcoma cells. *(B)* Imaging of tagged U2OS cell line showing endogenous SEC13-eGFP (in green) with cell nuclei visualized by DAPI staining (upper panel); imaging of endogenous SEC13 visualized with eGFP (green), showing its colocalization with ER tracker (in blue) (lower panel). (C) Immunofluorescence of U2OS cells showing proximity localization of endogenous eGFP-SEC13 (green) and endogenous UPF1 (red) (upper panel). Lower panel shows the localization of endogenous eGFP-UPF2 (green) and endogenous SEC13 detected with an anti-SEC13 antibody. In both cases, cell nuclei were visualized by DAPI staining. (*D, E*) Proximity ligation assay (PLA) using antibodies against eGFP alongside endogenous,UPF1 and UPF2 proteins generated a discrete signal (red spots), indicating that the proteins are <40 nm apart. *(D)* SEC13 is localized in close proximity of UPF1. *(E)* SEC13 is localized in close proximity of UPF2. The graphs in left panels D and E, show the quantification of PLA signal. Each point represents mean PLA count per nucleus in one captured frame. Significance was determined by two-tailed t-test.

First, we confirmed that tagged-SEC13 localizes to the endoplasmic reticulum (ER), as shown by colocalization of endogenous SEC13-eGFP with the ER tracker, as expected (Shaywitz et al. 1995)(Fig. 2B). In agreement, tagged-SEC13 is also associated with the translocon, as evidenced by its colocalization with SEC61B (Hartmann et al. 1994) (Supplemental Fig. S2C). We next investigated the localization of SEC13 in reference to two core NMD factors, UPF1 and UPF2. It has been reported by us and others, that UPF1 mainly localizes diffusely in the cytoplasm (Applequist et al. 1997; Serin et al. 2001; Longman et al. 2020), but was also found associated with ER-bound polysomes (Jagannathan et al. 2014). We had also previously shown that a fraction of UPF1 is anchored at the ER membrane, which was only evident following digitonin permeabilization, and interacts with the translocon, as revealed by proximity Ligation Assay (PLA) using antibodies against endogenous UPF1 and SEC61B proteins (Longman et al. 2020). Here, we observed that endogenous UPF1 localizes mainly to the cytoplasm, as expected, under these experimental conditions, and shows similar localization but little direct overlap with SEC13 (Fig. 2C, upper panel).

We next introduced an FKBP12^F36V^/3xFLAG/eGFP-tag in the *UPF2* locus in U2OS cells (U2OS-UPF2-tag), (Fig. 2A, lower panel), and imaged these cells, which showed the colocalization of endogenous UPF2-eGFP with endogenous SEC13 at the ER (Fig. 2C, lower panel). This agrees with our previous finding that a fraction of UPF2 localizes to the translocon in HeLa cells (Longman et al. 2020). Proximity Ligation Assay (PLA) detected a robust signal between UPF1 and SEC13 (GFP) in U2OS-SEC13-tagged cells (Fig. 2D), and between SEC13 (GFP) and UPF2 (Fig. 2E), confirming the proximity of SEC13 with both NMD factors. A PLA signal between SEC13 and SEC61B confirmed its association with the translocon (Supplemental Fig. S2D). Taken together these data demonstrates that SEC13 localizes to the ER in close proximity to core NMD factors. Combined with the AlphaFold structural data shown above, these data support a model in which SEC13 interacts directly with UPF2 thereby recruiting UPF1, UPF3 and SMG7 to the NMD complex.

### Activity of SEC13 in the NMD pathway

Fluorescence- and luminescence-based reporters are important tools for quantifying nonsense-mediated mRNA decay (NMD) activity, since they allow for rapid and sensitive measurements of NMD efficiency at the single-cell level. To quantify NMD activity at single-cell level, we developed a novel dual-color NMD sensor based on a series of previous NMD reporters (Pereverzev et al. 2015; Paillusson et al. 2005; Kolakada et al. 2024). This novel NMD sensor comprises a red fluorescent protein (mScarlet3) (Gadella et al. 2023), fused to downstream β-globin exons harboring an intron, such that the mScarlet3 stop codon acts as a premature termination codon (PTC) and renders the transcript NMD-sensitive, facilitating a quantitative fluorescent read-out of NMD activity at single-cell level. A second transcriptional unit, driven by the same human Ubiquitin C promoter, expresses TagGFP2 as a normalization control for reporter expression (NMD-sensor). This NMD sensor was inserted by CRISPR/Cas9 genome editing into the AAVS1 locus of WT U2OS cells (U2OS-NMD-sensor) (Fig. 3A). As expected, active NMD kept mScarlet3 levels low relative to TagGFP2 expression (Supplemental Fig. S3A, DMSO panel). In contrast, treatment with the potent NMD inhibitor SMG1i (Gopalsamy et al. 2012) drove a substantial increase in mScarlet3 without affecting TagGFP2 (Supplemental Fig. S3A, SMG1i panel). Next, we show that siRNA-mediated knock-down of UPF1, UPF2, SEC13 and NBAS (KD efficiencies shown in Supplemental Fig. S3B), resulted in NMD inhibition to varying degrees. UPF1 depletion had the most pronounced effect, with significant increases in the mScarlet3 geometric mean in all other KD conditions (Figs. 3A, lower panel, and Supplemental Fig. S3C). This was confirmed by quantification of mScarlet3 mRNA by qPCR (Supplemental Fig. S3D). To further confirm the role of SEC13 in the NMD pathway, we stably transduced U2OS-SEC13-tag cells with lentiviral NMD sensor and enriched for a polyclonal mScarlet3+ population (Fig. 3B).

**FIGURE 3.**
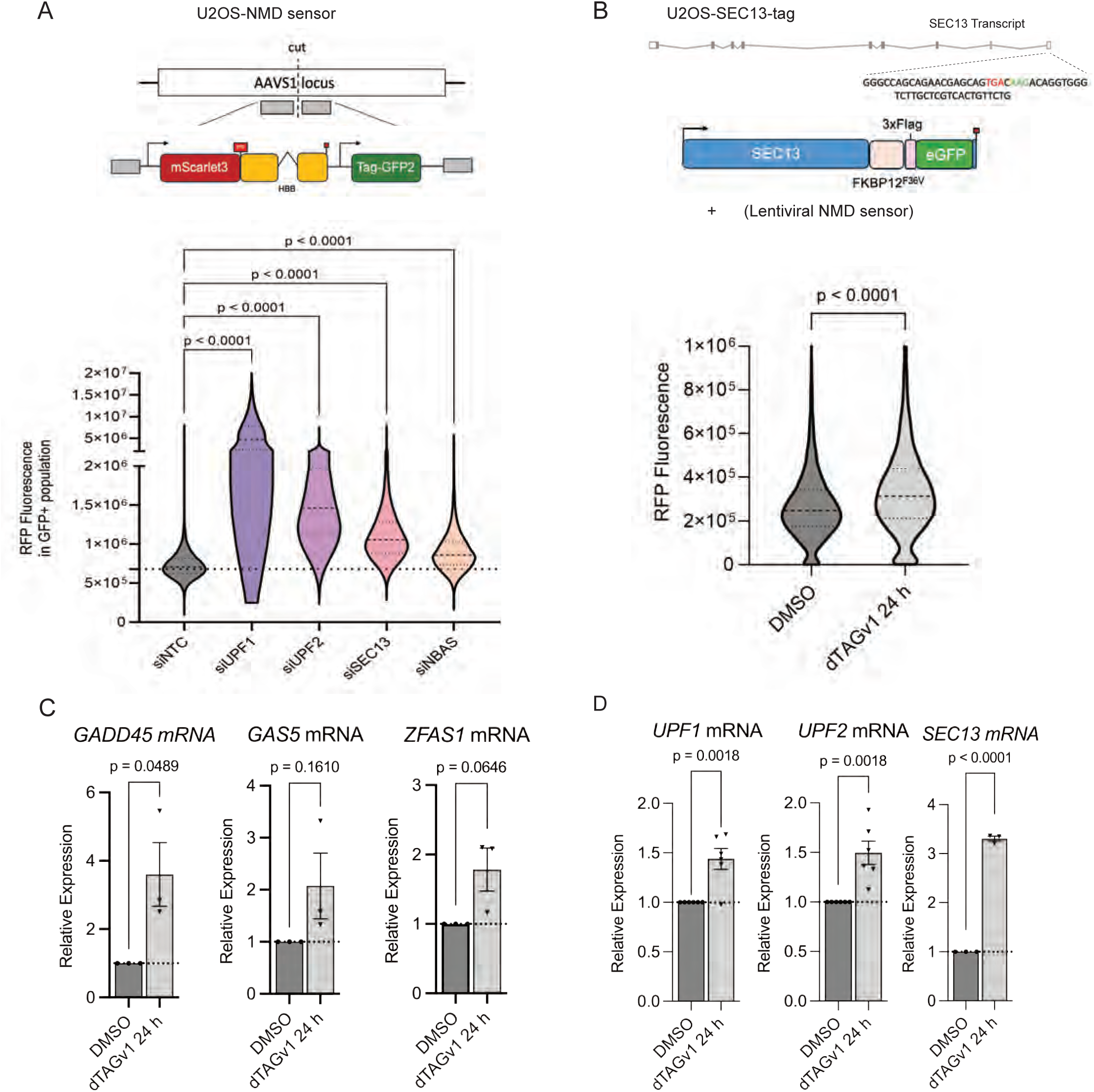
SEC13 is a bona fide NMD factor. *(A)* Schematic representation of a novel fluorescent NMD sensor in U2OS cells, comprising an N-terminal mScarlet3 sequence fused to downstream β-globin (HBB) exons harboring an intron, and a second TagGFP2 transcriptional control unit, that were inserted into the AAVS1 locus by CRISPR/Cas9. siRNA-mediated knock-down (KD) of UPF1, UPF2, SEC13 and NBAS, results in a shift in mScarlet3 expression measured by flow cytometry (n=5 biological replicates), as shown in a Violin plot; significance determined by one-way ANOVA with multiple-comparisons; p-value displayed. (*B*) Schematic representation of U2OS-SEC13-Tag; comprising a C-terminal FKBP12F36V, 3xFlag and eGFP inserted at the endogenous SEC13 locus. Lentiviral NMD-sensor transduction and bulk selection of mScarlet3+ cells; acute degradation of SEC13 following treatment with 300nM dTAGv1 for 24 h leads to mScarlet3 upregulation. mScarlet3 geometric mean quantified following dTAGv1 treatment (24 h); significance determined by unpaired t-test; p-value displayed. (*C*) Acute degradation of SEC13 following treatment with 300nM dTAGv1 for 24 h leads to upregulation of mRNAs of known NMD targets measured by qRT-PCR and normalized to POLR2J expression. Each point represents one biological replicate; bars represent mean with SEM. Significance determined by two tailed unpaired t-test. (*D*) Acute degradation of SEC13 following treatment with 300nM dTAGv1 for 24 h leads to upregulation of mRNAs encoding NMD factors, as measured by qRT-PCR and normalized to POLR2J expression. Each point represents one biological replicate; bars represent mean with SEM. Significance determined by two tailed unpaired t-test.

We acutely depleted SEC13 using dTAGv1 for 24 h and confirmed a significant increase in mScarlet3 geometric mean (Fig. 3B). We could clearly demonstrate that upon acute SEC13 degradation, the steady-state levels of three well-characterized NMD targets, Zfas1, Gadd45, and Gas 5 (Boehm et al. 2025), were significantly upregulated, as measured by qRT-PCR (Fig. 3C). Interestingly, acute SEC13 degradation also led to the upregulation of mRNAs encoding core NMD factors, UPF1 and UPF2 and SEC13 (Fig. 3D), consistent with the previously described negative feedback loop mechanism, whereby NMD controls the levels of transcripts encoding NMD factors (Huang et al. 2011; Yepiskoposyan et al. 2011; Longman et al. 2013). Taken together, these results clearly demonstrate that SEC13 is a bona fide NMD factor.

### SEC13 regulates a subset of NMD targets translated both at the cytoplasm and at the ER

To identify RNA targets that are regulated by SEC13 genome-wide, we profiled mRNA abundance changes by RNA-sequencing following SEC13 depletion and compared these with UPF2 knockdown and a previously published UPF1 knockdown dataset (Longman et al. 2020). Knockdown efficiency was confirmed by qRT-PCR (Supplemental Fig. S4A), in agreement with reduction measured by RNA sequencing (SEC13: fold change −1.90, padj = 4.5 × 10^−65^; UPF2: fold change −2.32, padj = 3.01 × 10^−99^). Depletion of SEC13 significantly affected the mRNAs of 4,633 genes, leading to an increase in the steady-state levels of 2,553 mRNAs (padj < 0.01; Fig. 4A; Supplemental Table S2). Depletion of UPF2 resulted in expression changes in 3,908 genes, with increased expression of 2,253 genes (Supplemental Fig. S4B; Supplemental Table S3). Reactome pathway analysis of SEC13-upregulated genes (padj < 0.01) revealed strong enrichment for ER stress-related pathways (Fig. 4B) (Ragueneau et al. 2026), consistent with a role for SEC13 at the ER. Top enriched pathways included XBP1-mediated activation of chaperone genes (p < 3.46 × 10^−5^), IRE1α signaling (p < 4.34 × 10^−5^ and the unfolded protein response (UPR; p < 2.13 × 10^−5^), suggesting a role for SEC13-dependent NMD in the regulation of ER stress. Next, we classified genes commonly upregulated by both UPF2 and UPF1 depletion (Longman et al. 2020), as high confidence NMD targets. We observed co-regulation of mRNAs when UPF2, UPF1 or SEC13 were depleted with 164 common targets across all three knockdowns (Fig. 4C), indicating that SEC13 co-regulates a core subset of canonical NMD targets. Depletion of SEC13 and UPF2 produced positively correlated transcriptome-wide expression changes (R = 0.768, p < 1.6 × 10^−225^; Fig. 4D), confirming that both factors regulate a common set of targets.

**FIGURE 4.**
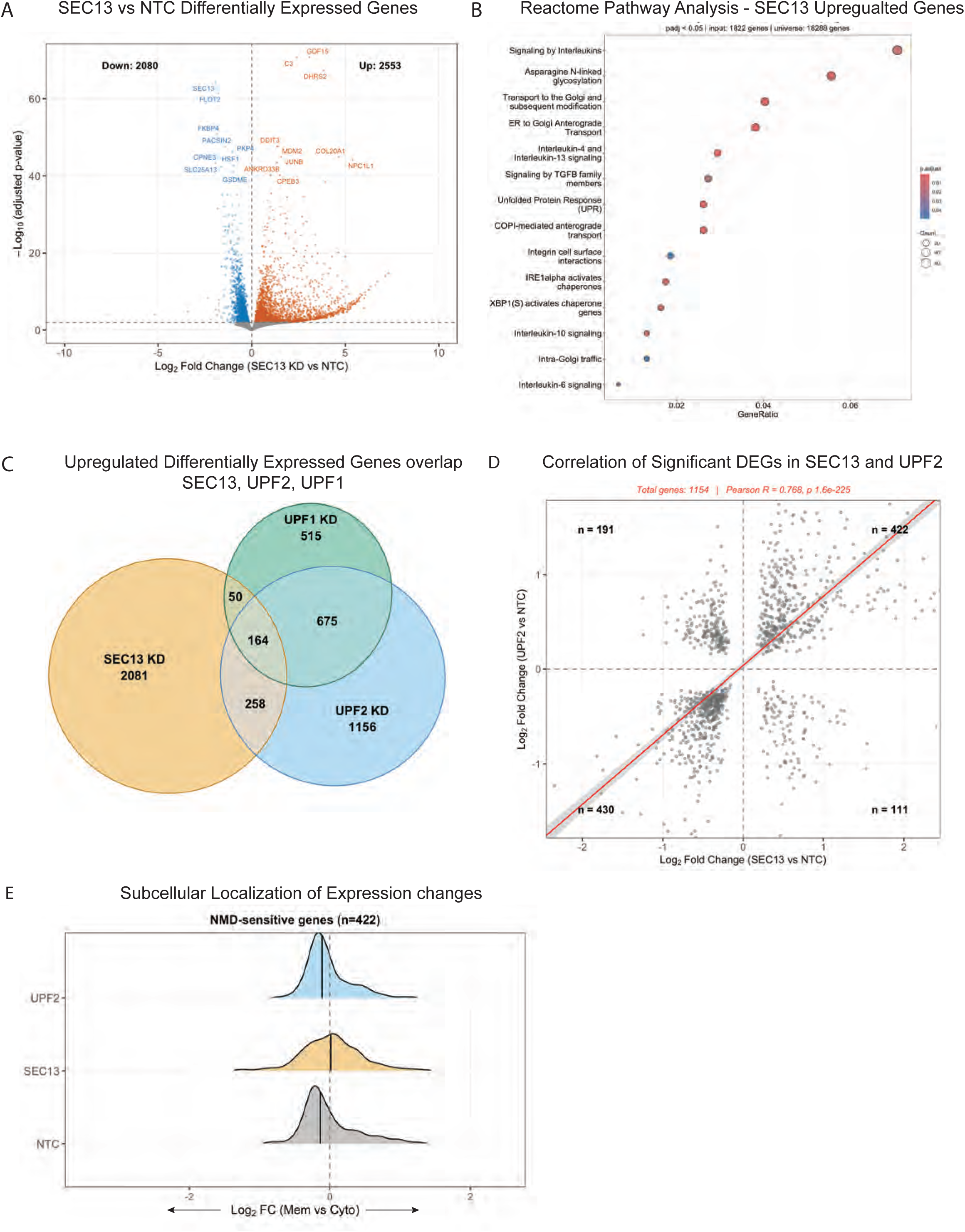
SEC13 depletion reveals common NMD targets with UPF1 and UPF2 and implicates SEC13 in the regulation of ER stress. *(A)* Volcano plot showing differentially expressed genes upon siRNA-mediated depletion of SEC13; genes with LFC > 1 and padj < 0.001 are indicated, with gene numbers shown in brackets. *(B)* Reactome pathway analysis of all significantly upregulated genes upon SEC13 depletion, identifying enrichment for unfolded protein response pathway. *(C)* Venn diagram showing overlap of significantly upregulated genes following siRNA knockdown of SEC13, UPF2 (this study) and UPF1 (Longman et al. 2020). (*D*) Correlation of log₂ fold change upon SEC13 vs UPF2 depletion; Pearson correlation R = 0.77; experimentally validated ER-associated genes highlighted in blue. *(E)* Log_2_FC of SEC13 and UPF2 common NMD Targets (422 genes); on comparison of membrane and cytoplasmic fractionation samples (Log2FC Mem vs Cyto) showing shift in expression distribution upon KD. Positive number expression is higher in membrane samples; negative expression is higher in cytoplasmic samples.

To resolve the subcellular distribution of SEC13 NMD targets, we performed RNA-seq on cytoplasmic and membrane fractions from SEC13- and UPF2-depleted HeLa cells. Fractionation efficiency was validated by the expected compartment-specific distribution of marker proteins and mRNAs (Supplemental Figs. S4C, D). We examined the subcellular distribution of NMD targets that were commonly upregulated upon depletion of both UPF2 and SEC13. Specifically, we compared transcript levels between the membrane and cytoplasm (log₂ fold change, membrane vs. cytoplasm) in UPF2-depleted, SEC13-depleted, and mock-depleted (NTC) cells. NMD targets showed predominantly cytoplasmic distribution in both UPF2-depleted and mock-depleted (NTC) cells. In contrast, depletion of SEC13 resulted in a modest shift of NMD targets toward the membrane fraction compared to NTC, suggesting SEC13 is more significantly influencing the expression of these targets at the membrane (Fig. 4E). Taken together, these data demonstrate that SEC13 participates in both cytoplasmic and ER-localized NMD, with the capacity to regulate a broad range of targets.

### Separation-of-function of SEC13 in NMD and other cellular complexes

To determine whether the role of SEC13 in NMD operates independently of its functions in the COPII, nuclear pore (NPC) or GATOR2 complexes, we performed knockdown of representative components of each pathway, including SEC23 (COPII), NUP133 (NPC), SEH1L (NPC and GATOR2) and WDR24 (GATOR2), alongside UPF2 as a positive NMD control. Knockdown efficiencies were confirmed by qRT-PCR (Supplemental Fig. S5A). Using the U2OS NMD sensor to assess NMD activity, depletion of SEH1L, WDR24 and SEC31 had little effect on mScarlet3 fluorescence, initially suggesting that the NMD activity of SEC13 operates independently of its roles in the GATOR2 complex, NPC, or COPII (Fig. 5A). By contrast knockdown of SEC23, a core COPII component produced increased fluorescent signal of similar magnitude to SEC13 (Fig. 5A). A confounding factor in interpreting these results is that depletion of any single component of a multi-protein complex can potentially perturb the levels of its binding partners, making a clear functional separation difficult. For instance, SEC23 knockdown also reduced SEC13 protein levels while simultaneously increasing SEC13 mRNA (Supplemental Figs. S5B, C). This is consistent with reduced SEC13 protein stability in the absence of its COPII binding partner(s) and a compensatory transcriptional response. Furthermore, acute SEC13 depletion upregulated SEC23 mRNA level (Supplemental Fig. S5D), reinforcing a mutual interdependence between these proteins. Therefore, we cannot distinguish between the direct NMD role for SEC23 itself, or its effect on NMD as a consequence of a secondary SEC13 loss. In a similar fashion, knockdown of NUP133, an early assembly factor of the NPC outer ring, also produced a positive NMD sensor signal (Fig. 5A). Similarly to SEC23, it also reduced SEC13 protein level (Supplemental Fig. S5B), whilst upregulating SEC13 mRNA (Supplemental Fig 5C), and acute SEC13 depletion downregulated NUP133 mRNA (Supplemental Fig. S5D). From these experiments, we can conclude that the role of SEC13 in NMD operates independently of its function in the GATOR2 complex. By contrast, the results observed with COPII and NPC components could be due to indirect effects of these components in regulating SEC13 level, and therefore we cannot rule out at this stage some contribution of these pathways to NMD using this dataset. Within the NPC, the lack of effect of SEH1L depletion could be rationalized by its position downstream of SEC13 in outer ring assembly, therefore having limited effect on SEC13 protein or mRNA levels. An interaction between Upf1p and the nuclear pore (Nup) proteins, Nup100p and Nup116p has been shown in *S. cerevisiae* (Nazarenus et al. 2005), but there is not clear indication that the NPC is involved in the NMD response.

**FIGURE 5.**
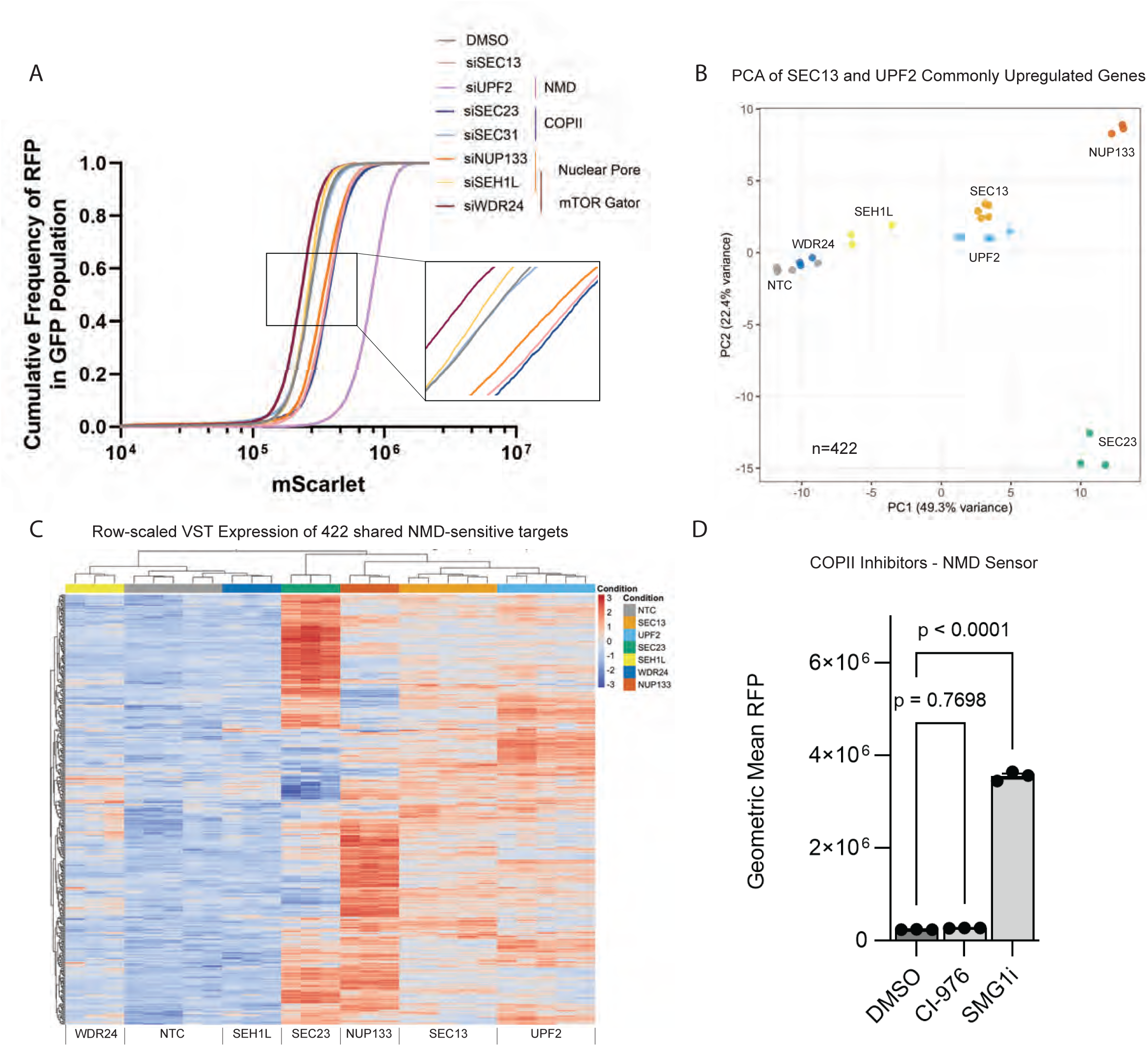
Dissecting the NMD function of SEC13 from its roles in other cellular complexes. *(A)* Cumulative mScarlet3 fluorescence measured by flow cytometry following siRNA knockdown of UPF2 (NMD control), SEC13, NPC components (NUP133, SEH1L), GATOR2 components (WDR24, SEH1L) and COPII components (SEC23, SEC31). Shift in mScarlet3 signal indicates NMD inhibition. Zoomed-in panel to highlight central overlap of samples. *(B)* PCA of genes commonly upregulated by SEC13 and UPF2 knockdown (SEC13 NMD target set), showing separation of SEC13 and UPF2 from SEC23, NUP133 and NTC. *(C)* Heatmap of SEC13 NMD targets showing a differential expression pattern for these genes in comparison with UPF2; highlighting a distinct but not unique role of SEC13 in NMD, COPII trafficking and nuclear pore formation. (D) 24 h treatment with COPII inhibitor CI-976 show non-significant changes in mScarlet florescence as measured by Flow Cytometry.

To further resolve whether SEC13 regulates NMD independently of its roles in other complexes, we performed RNA-seq following knockdown of SEC13, UPF2, SEC23, NUP133, SEH1L and WDR24 (Supplemental Fig. S5E). Principal component analysis (PCA) of the top 1,000 differentially expressed genes separated conditions broadly across PC1 and PC2 (Supplemental Fig. S5F). Next, we selected genes upregulated by both SEC13 and UPF2 knockdown as a high-confidence SEC13 NMD targets. When the analysis was restricted to this set of SEC13 NMD targets, SEC13 and UPF2 were clearly separated from SEC23, NUP133 and the control (NTC) (Fig. 5B). Hierarchical clustering of these shared targets revealed a distinct expression signature for SEC13 and UPF2 that was not recapitulated by knockdown of COPII or NPC components (Fig. 5C), suggesting that the impact of SEC13 on NMD targets most likely acts independently of its roles in the COPII and the NPC complexes. This was further evidenced by treatment of U2OS -NMD sensor cells with an inhibitor of COPII function, C1-976 (Brown et al. 2008), known to reduce the COPII late-stage vesicle budding after complete formation. This treatment did not lead to NMD target upregulation, indicating that COPII-mediated ER export is not required for NMD (Fig. 5D). To further support this separation of function at the structural level, AlphaFold modeling predicted a direct interaction between SEC13 and UPF2 (Fig. 1D) yet showed no interaction between UPF2 and components of the COPII, NPC or GATOR2 pathways (Supplemental Fig. S5G). Taken together, these results support a bona fide role for SEC13 in NMD that operates in both the cytoplasm and at the ER, and that is likely distinct from its other cellular functions.

### NMD and the unfolded protein response

The Unfolded Protein Response (UPR) is activated by ER stress arising from the accumulation of misfolded proteins in the ER (Walter and Ron 2011; Acosta-Alvear et al. 2025). Since chronic activation of the UPR contributes to many human pathologies, the fidelity of UPR activation must be tightly regulated. Previous evidence highlighted a role for the NMD pathway in the modulation of the ER stress response, by ensuring appropriate activation of the UPR (Goetz and Wilkinson 2017; Karam et al. 2015; Sieber et al. 2016). It has also been shown that mRNAs encoding sensors of the UPR are targeted by NMD, including IRE1α, as well as ATF-4 and CHOP that are activated by PERK branch signaling. Since SEC13 is an ER-localized NMD factor, we wanted to confirm and extend these observations, and asses the role of SEC13 in NMD-mediated regulation of the UPR.

First, we used U2OS-SEC13-tag cells to induce degron-mediated acute degradation of SEC13 and observed upregulation of mRNAs encoding sensors for all three branches of UPR, namely *IRE1, ATF6* and *PERK* (Fig. 6A). To broaden this analysis, we compared the effects of acute SEC13 depletion with siRNA knockdown of an ER-NMD factor NBAS, and dTAG degradation of the core NMD factors UPF1 (UPF1-FKBP12^F36V^ 3xFlag eGFP) and UPF2 on a panel of mRNAs encoding ER stress components across multiple tagged cell lines (Supplemental Fig. S6A). We observed that SEC13 regulates the stability of ATF4 and CHOP; which is also affected by KD of UPF1, UPF2 and NBAS. Thus, SEC13 seems to control the stability of ER stress sensors of all branches of the UPR, in line with reported effects for core NMD factors (Fig. 6A and Supplemental Fig. S6A)(Karam et al. 2015).

**FIGURE 6.**
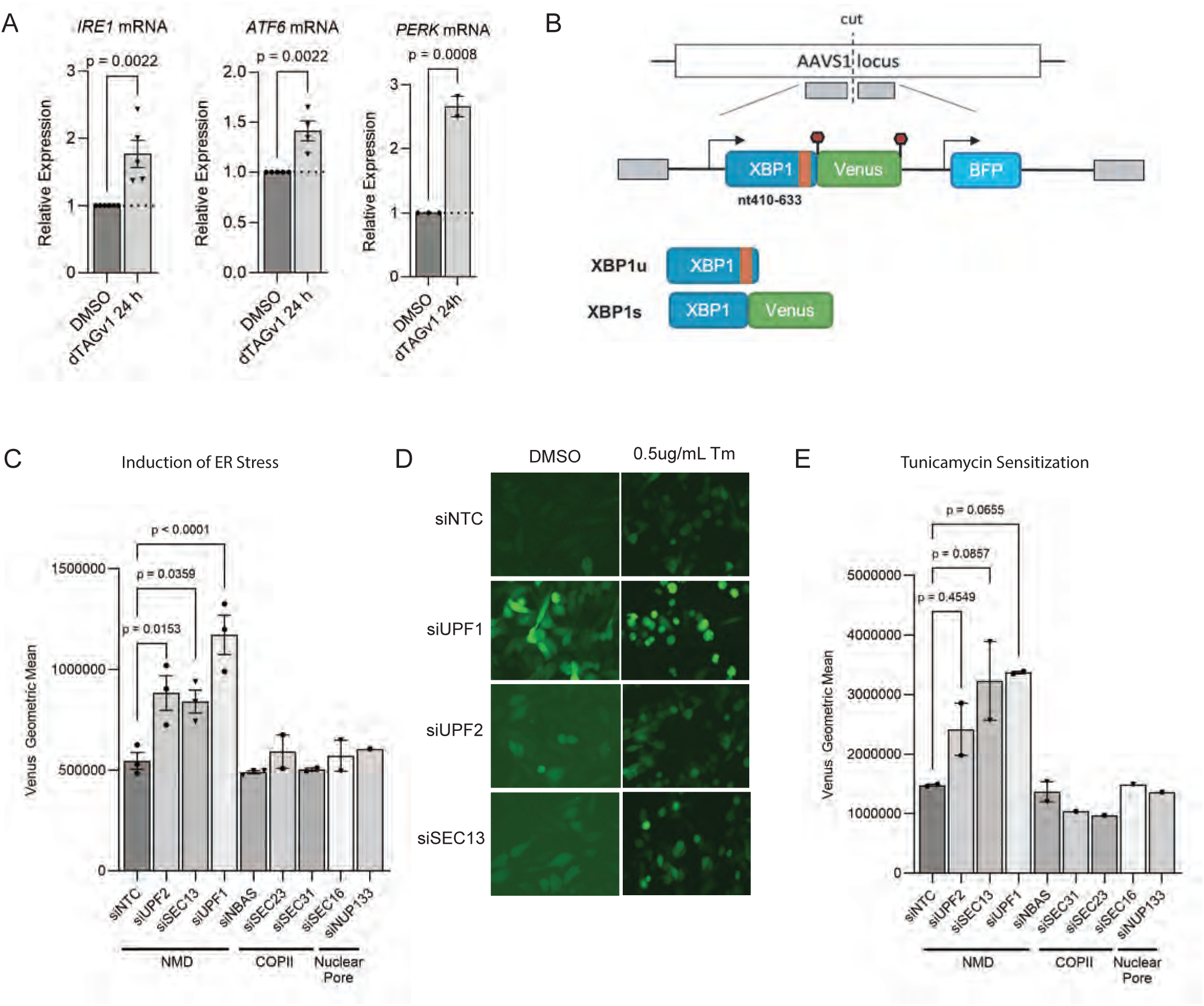
Disruption of the NMD pathway induces ER stress and sensitizes to the effect of tunicamycin. *(A*) Steady-state levels of mRNAs encoding ER stress sensors from the IRE1 (Ire1α), ATF6 (Atf6) and PERK branches, following dTAGv1-induced degradation of SEC13 for 24 h. Quantitation was done by qRT-PCR and normalized to POLR2J. Each point represents one biological replicate; bars represent mean with Standard error of the mean (SEM). Significance determined by two tailed t-test. (*B*) Schematic of the ERAI ER stress reporter inserted into the AAVS1 locus in U2OS cells. XBP1 and Venus coding regions are indicated by blue and green boxes, respectively; the non-canonical intron is shown in red. A second transcriptional unit drives BFP expression as a control. XBP1u: unspliced (stop codon precludes Venus expression); XBP1s: IRE1-spliced (Venus expressed). (*C*) Venus fluorescence measured by flow cytometry following siRNA knockdown of NMD factors and components of the COPII and nuclear pore complexes. Geometric mean of all events; each point represents one biological replicate; bars represent mean ± SEM. Significance determined by one-way ANOVA with multiple comparisons; p-values displayed. *(D*) Representative imaging of ERAI sensor cells treated with tunicamycin (0.5 µg/mL, 24 h) or DMSO, following siRNA knockdown of NMD factors. (*E*) Venus fluorescence measured by flow cytometry following siRNA knockdown and incubation with a single concentration of tunicamycin (2 µg/mL, 24 h). Geometric mean of all events; each point represents one biological replicate; bars represent mean ± SEM. Significance determined by one-way ANOVA with multiple comparisons; p-values displayed.

To monitor ER stress in mammalian cells in culture, we adapted a previously described reporter, termed ‘ER stress–activated indicator’ (ERAI) that monitors the non-conventional splicing of the mRNA encoding the transcription factor XBP1 (Iwawaki et al. 2004). In this reporter, the XBP1 sequence is fused to Venus, a variant of green fluorescent protein, such that upon ER stress induction, IRE-1α is activated and eliminates this non-conventional 26nt intron, thus leading to an XBP-1-Venus fusion protein, which can be detected by its fluorescence. We modified the original sensor (see methods), and we inserted this reporter into the AAVS1 locus of U2OS osteosarcoma cells, using CRISPR-Cas9 genome editing, followed by single cell sorting to obtain a clonal ER stress-responsive population (Fig. 6B). Treatment with tunicamycin (Tm), a potent inhibitor of protein glycosylation induces ER stress, resulted in a large increase in Venus fluorescence, validating the reporter. Importantly, this effect was partially abrogated by STF-083-010 a selective inhibitor of IRE1α endonuclease activity (Papandreou et al. 2011) (Supplemental Fig. S6B) To broadly investigate the role of the NMD pathway in ER stress, we monitor the ER stress sensor reporter, upon knock-down of individual NMD factors. Importantly, knock-down of UPF1, UPF2 and SEC13 leads to induction of ER stress, as shown by increased Venus fluorescence (Fig. 6C), and confirmed by representative imaging (Fig. 6D). Importantly, in the case of SEC13, this effect seems to be NMD-specific and to operate independently of the role of SEC13 in other cellular complexes, since knock-down of COPII components (SEC16, SEC23 and SEC31) or nuclear pore components (NUP133) did not result in an induction of ER stress (Fig. 6C). Furthermore, when corrected for the effect of siRNA knockdown alone on ER stress, depletion of NMD factors increased sensitivity to tunicamycin-induced ER stress, (Fig. 6E), in a dose-responsive manner (Supplemental Fig. S6C).

## DISCUSSION

We initially identified SEC13 as a novel NMD factor in a functional genome-wide RNAi screen in *C. elegans* and showed that it can regulate NMD in human cells (Casadio et al. 2015). However, the mechanistic basis for this multifunctional protein in the NMD pathway remained unexplored. Here, we used proteomics, IP-Western analysis and PLA assays to demonstrate the interaction of SEC13 with core NMD factors, including UPF1, UPF2, UPF3 and SMG7. AlphaFold structural modeling suggested that UPF2 interacts directly with SEC13, which could bring the other NMD factors to an NMD complex containing SEC13 (Figs. 1 and 2). We developed a novel highly sensitive fluorescent NMD sensor that clearly shows the effect of SEC13 on NMD, with a comparable magnitude of NMD sensor upregulation to core NMD factors, UPF1 and UPF2. More importantly SEC13 also regulates the stability of well-characterized endogenous NMD targets and participates in the feed-back loop regulating the steady-state levels of mRNAs encoding NMD factors (Fig.3).

The NMD pathway is intimately coupled to mRNA translation and regulates the stability of mRNAs translated by cytoplasmic ribosomes (Trcek et al. 2013). We previously discovered that the ER-localized NBAS, which is involved in retrograde transport from the Golgi-to-ER, has an independent function in NMD at the ER, preferentially targeting mRNAs involved in the cellular stress response (Longman et al. 2020). The ER is a site of localized protein synthesis, where translation of transmembrane and secreted proteins occurs on ER-bound ribosomes (Reid and Nicchitta 2015). Indeed, several RNA decay pathways at the ER, including regulated inositol-requiring enzyme 1α (IRE1α)-dependent mRNA decay (RIDD) (Hollien et al. 2009), and Argonaute-dependent RNA silencing (Efstathiou et al. 2022; Ottens et al. 2024), have been described. Our initial discovery that SEC13 localizes to the membrane of the ER, raised the possibility that this protein could also be involved in an ER-localized NMD pathway, in a similar manner to NBAS (Casadio et al. 2015; Longman et al. 2020). The functional characterization of SEC13 in the NMD pathway provided here show some similarities with NBAS, but also important differences. Whereas both proteins regulate the stability of mRNAs translated at the ER, NBAS seems to be dedicated preferentially to the ER-NMD pathway, most likely due to its intrinsic localization to the ER. By contrast, the moonlighting SEC13 protein, regulates the stability of mRNAs translated both in the cytoplasm and also at the ER, in agreement with its wide subcellular distribution. Thus, it would appear that the needs of cells to regulate NMD of mRNAs translated at the ER is facilitated by auxiliary NMD factors, such as NBAS, that primarily regulates ER-NMD, but also by factors, such as SEC13 that localize to the ER and to the nuclear pore, and regulate the stability of mRNAs translated both at the ER and in the cytoplasm. Several lines of evidence support a role for SEC13 in NMD regulation, operating independently from its function as a component of the GATOR2, COPII and NPC complexes (Fig. 5 and Supplemental Fig. 5).

There seem to be several links between the NMD pathway and the ER stress response (Goetz and Wilkinson 2017). A non-canonical UPF1 isoform, UPF1_LL,_, functions in NMD upon activation of the integrated stress response (ISR) resulting in repression of translation and induction of stress response genes, which are potential NMD targets (Fritz et al. 2022; Gowravaram et al. 2018). The previously proposed role for the NMD pathway in the regulation of the UPR could act through a dual mechanism. First, by degrading RNA targets that would otherwise give rise to truncated and/or misfolded proteins, thus decreasing the ER load, and limiting the UPR. Secondly, NMD affects the stability of UPR sensors, including IRE1α, ATF-4, and CHOP, thereby controlling the threshold of cellular stress necessary to activate the UPR (Goetz and Wilkinson 2017; Karam et al. 2015)). We confirmed and extended those observations and demonstrated an NMD-mediated role for SEC13 in UPR regulation by affecting the stability of ER stress sensors of the three branches of the UPR (Fig. 6 and Supplemental Fig. S6).

There are several examples of moonlighting functions of proteins involved in RNA processing, which include several glycolytic enzymes, such as Glyceraldehyde-3-phosphate dehydrogenase (GAPDH), that were found to bind to RNA and have been proposed as post-transcriptional regulatory roles (Castello et al. 2015; Wegener and Dietz 2022). Related to NMD, we showed that NBAS acts in ER-NMD, independently of its role in Golgi-ER retrograde transport (Longman et al. 2020). Here, we add SEC13 to this set of moonlighting proteins with a role in RNA regulation. In summary, we have here characterized a defined role for SEC13 in NMD regulation, that seems to operate independently of its other cellular functions, highlighting a moonlighting function of SEC13 in RNA regulation.

## MATERIALS AND METHODS

### Cell Culture and Transfections

HeLa and U2OS cells were maintained in DMEM media with high glucose, GlutaMAX™ Supplement, pyruvate (Gibco Life technologies; 10569010) supplemented with 10% FCS, at 37°C in the presence of 5% CO_2_. Tagged and sensor integrated cell lines were maintained in the same media. Transfections were carried out in Opti-MEM reduced serum medium (Gibco, 31985047). Transfections of siRNA oligos were done using either DharmaFECT 1 (Dharmacon, T-2001-03) or Lipofectamine 2000 (Thermo Fisher Scientific, 11668019) following manufacturer’s protocol. For NMD assays cells were plated in 6 well plates and transfected with 60pmol of indicated siRNAs; before a second transfection on the third day and harvesting 48hrs later. For total RNA-sequencing and subcellular fractionation; cells were plated in 6-well plates and transfected with 60pmol of indicated siRNAs. Cells were expanded into 10 cm plates the following day and were transfected with 120pmol of the same siRNAs on day 3 and were harvested for analysis 4 days after the first depletion.

### Design and screening of CRISPR cell lines

Guide RNAs (gRNAs) were designed using CHOPCHOP https://chopchop.cbu.uib.no/Cas-Designer. Guides were cloned into pSpCas9(BB)-2A-Puro (px459) V2.0 (Ran et al. 2013)(Ran et al. 2013). For the tagged cell line, four gRNAs were selected by closest proximity to the start codon and highest predicted efficiency. For each guide RNA, the complementary sequence was determined and a *Bbs*I restriction site added to both oligos. Designed gRNAs were ordered as custom single stranded DNA oligos with an extra G at the 5’ end, and 5’ phosphate at the reverse complement DNA strand (IDT). Top and bottom strands of gRNAs were annealed at a concentration of 100 μM and cloned into the px459 V2.0 vector using *Bbs*I restriction cloning. 1 μL of a 1:200 dilution of annealed gRNAs was ligated with the T4 DNA Ligase (NEB) into 36 ng of the px459 V2.0 vector. To assess cutting efficiency of gRNAs, each plasmid was transfected into U2OS cells and DNA extracted as above. PCR over the target region was performed with custom primers (Supplemental Table S4) and the resulting products assessed by DNA electrophoresis. PCR products were also sent for Sanger sequencing, and traces were analysed ICE to give exact cutting efficiencies for each guide. Repair template was ordered as a custom plasmid from IDT. The repair template containing synthetic homology arms, 3XFLAG-tag and eGFP with or without FKBP12^F36V^ with mutated PAM sites was cloned into the pGEM T-Easy backbone using TA Cloning. The gRNA/Cas9 plasmid and linearized repair template were transfected and selected with 1 µg/ml puromycin for 48 hours. 5 days post-transfection surviving cells were sorted by Fluorescence-Activated Cell Sorting (FACS) using the Cytoflex SRT system into 96 well plates and expanded. Colonies were PCR screened and correct targeting verified by Sanger sequencing. Sequences sgRNAs are listed in Supplemental Table S4. For integration of NMD Sensor and ERAI; Assemblies were performed using NEBuilder® HiFi DNA Assembly Master Mix (NEB). For the ERAI, important components of the sensor were cloned individually and assembled using HiFi technology. To create stable isogenic cell lines, donor plasmids were constructed for CRISPR-mediated integration into the human AAVS1 safe-harbour locus. First, ∼400 bp homology arms flanking the target site were amplified from genomic DNA isolated from U2OS cells. The desired NMD-sensor or ERAI cassettes were then assembled between these homology arms into a pcDNA3.1 backbone using HiFi DNA Assembly, creating the final pAAVS1 donor plasmids. U2OS cells were co-transfected with pX459-gAAVS1 plasmid and the respective donor plasmid using Lipofectamine 3000 (Thermo Fisher Scientific). Twenty-four hours post-transfection, cells were selected with 2 µg/mL Puromycin for 48 hours. The surviving cell population was expanded for approximately one week before a pure population of reporter-expressing cells was isolated by fluorescence-activated cell sorting (FACS) into 96-well plates. Clones were expanded and validated by genomic PCR to confirm correct integration.

### Immunofluorescence

Cells were grown on coverslips, fixed with 4% paraformaldehyde at room temperature for 10 min, washed with PBS and permeabilized with 0.2% Triton X-100 at room temperature for 10 min. Coverslips were then incubated for 1 h with block buffer (1% BSA, 0.01% Triton X-100 in PBS), followed by primary antibodies (diluted 1:200-500 in block buffer) in a humidified chamber overnight at 4°C. Coverslips were washed 3 times with wash buffer (0.01% Triton X-100 in PBS). Secondary antibodies (Alexa Flour 488 or Alexa Flour 594, Molecular Probes, diluted 1:1000 in block buffer) were incubated with coverslips in a dark, humidified chamber for 1 h at room temperature. Coverslips were then washed 3 times with wash buffer and stained with 4,6-diaminidino-2-phenylidole (DAPI) at 50ng/ml, mounted in Vectashield (Vector) and sealed with nail varnish.

### Proximity ligation assay

Proximity ligation assay (PLA) PLA assay was performed following Duolink PLA fluorescence protocol (Sigma-Aldrich). Cells were grown, fixed, and permeabilized as for immunofluorescence on 8 well ibidi chamber slides. Slides were then incubated for 1 h with 1% BSA, 0.01% Triton X-100 in PBS block buffer provided with the Duolink in situ PLA probes at room temperature and incubated with primary antibodies. Slides were then washed three times with Duolink wash buffer A and Duolink probe incubation, ligation, and PLA signal amplification were performed using Duolink in situ detection reagent red kit according to the manufacturer’s instructions. In the last wash, coverslips were incubated with DAPI at 50 ng/ mL in 0.01× wash buffer B for 5 min and maintained in fresh PBS at 4°C for imaging.

### Image Capture and analysis

For immunofluorescence experiments images were acquired Epifluorescent images were acquired using a Hamamatsu Orca Flash 4.0 sCMOS camera (Hamamatsu, Japan) and a Zeiss Axio Imager.A1 fluorescence microscope with Plan-Neofluar/Apochromat objective lenses (Carl Zeiss, Cambridge, UK), an X-Cite Xylis LED light source (Excelitas Technologies) and Chroma #83000 triple band pass filter set (Chroma Technology Corp, Rockingham, VT, USA) with the single excitation and emission filters installed in motorised filter wheels (Prior Scientific Instruments, Cambridge, UK). Image capture was performed using Micromanager (Version 2.0). For PLA experiments images were acquired on a Nikon AX confocal microscope (Nikon Europe B.V.) using a 63X oil objective. The microscope comprises of a Nikon Eclipse Ti2 inverted microscope with Perfect Focus System and is equipped with a LUA-S6 laser bed (405, 458, 488, 514, 640 diode lasers). Detection is via four Photomultiplier tubes (2x Multi-Alkali Photomultiplier tubes and 2x GaAsP PMTs). Data were acquired using NIS Elements AR software (Nikon Europe B.V.). Z-stacks of images were acquired with a 0.4 μm step, scan size 2048x2048, 1.44x zoom and 2x frame averaging. Image analysis was carried out using the FIJI/ImageJ software. Automated image analysis was performed using a custom macro developed in Fiji (Schindelin et al. 2012). Briefly, 3D image stacks were converted to maximum intensity projections. Nuclei were segmented using Otsu thresholding on the DAPI channel and cell outlines identified. PLA spots were identified and quantified using a difference of gaussians method followed by local maxima detection on the Texas Red (624nm) channel. Statistics are reported as the total number of spots in a field per cell in that field of view. PLA analysis was carried out using a locally coded Image J Macro.

### Immunoprecipitation Western Blotting

Cells were washed and harvested in ice-cold PBS before pellets were lysed with immunoprecipitation (IP) buffer (20 mM Tris-HCl pH 8, 150 mM NaCl, 1mM EDTA, 1% NP-40, 0.2% Deoxycholate, Complete Protease Inhibitor (Roche), Phospho STOP (Roche), 1 mM DTT) for 20 min on ice. Anti-GFP MA (Chromotek) magnetic beads were washed, bound proteins were eluted with NuPAGE LDS sample buffer supplemented with reducing agent (Thermo Fisher). Proteins were resolved by SDS-PAGE on NuPAGE 4-12% Bis Tris or 3-8% Tris-Acetate precast gels (Thermo Fisher) and protein transfer was achieved using the iBlot™ 2 Gel Horizontal Transfer Device (Thermo Fisher) or wet transfer with 20% methanol 1x Transfer buffer at 15v for 60-90 mins. Nitrocellulose membranes were blocked in 5% BSA in PBS/Tween 20 (0.1%) and probed with the appropriate primary antibody diluted in blocking solution 1:1000. HRP-conjugated secondary antibodies (BioRad) were used at 1:10,000 and blots developed with ChemiGlow detection reagent and visualized using Image Quant LAS 4000 or 800 chemiluminescent camera.

### Immunoprecipitation Mass Spectrometry

Cells were harvested and lysed as in immunoprecipitation protocol (above). α-GFP antibody-coupled magnetic beads (Chromotek) were equilibrated with IP buffer. Lysates were resuspended in 500 μL IP buffer for capture of SEC13-FLAG-GFP bound proteins and subsequent mass spectrometry analysis. Immunoprecipitation were performed on Kingfisher Duo robot (Thermo Fisher) and subjected to in solution digestion using Trypsin (Thermo Fisher) according to standard protocols. for 4 h. Fractionated peptides were separated and analysed using a Dionex RSLC Nano system coupled to a Thermo Q-Exactive Plus instrument. (Thermo Fisher Scientific). Raw MS data were analysed using MaxQuant (v 1.5.6.5) (Max Planck Institute of Biochemistry) in conjunction with UniProt human reference proteome release 2016\_11 (uniprot.com), with match between runs (MS/MS not required), LFQ with 1 peptide required, and statistical analyses performed in R (RStudio 1.1.453 / R x64 3.4.4) (rstudio.com) using Wasim Aftab’s LIMMA Pipeline Proteomics (github.com/wasimaftab/LIMMA-pipeline-proteomics) implementing a Bayes-moderated method. Interactome analysis including gene ontology was carried out by inputting protein list into STRING (string-db.org/) and Gene Ontology enRIchment anaLysis and visuaLizAtion (GOrilla) (http://cbl-gorilla.cs.technion.ac.il/) (Eden et al. 2009).

### AlphaFold

AlphaFold3 (AlphaFoldServer) (Abramson et al. 2024) was run to predict pairwise interactions between full-length SEC13 and various known binding partners and NMD factors. Default parameters were used. To analyse the predictions produced by AlphaFold we used an analysis pipeline (Predictome) available online developed by the Walter laboratory (https://predictomes.org/).

### siRNA KD

NMD reporter U2OS cells stably expressing NMD sensor III or HeLa cells were mock-depleted or depleted twice of UPF1, UPF2, SEC13, NBAS, SEC23, SEC31, SEH1L, WDR24 and NUP133, and harvested 3-6 d after the first depletion and cells collected for Flow Cytometry and RNA analysis. NMD activity was determined by quantitative measurement of red and green fluorescence using Cytoflex flow cytometry analyser.

### Flow Cytometry

Following appropriate treatment cells were trypsinized, washed with cold PBS and placed on ice until analysis. Analysis was carried out using Cytoflex flow cytometry analyser. Gates were set using a nonfluorescent control. Green fluorescence expression (Blue-525) was analyzed by gating the intact cell population in FSC-A/ SSC-A scatter plot, followed by single cell exclusion gate in FSC-A/ FSC-H scatter plot, followed by gating only the red fluorescent-positive cells in YG-585. Gate settings were kept constant during the experiment. Data were analyzed with FlowJo software (version 10.6.0).

### dTAG treatment

Cells were treated for varying timepoints with 300nM dTAGv1 ligand and the GFP expression quantified by flow cytometry and protein expression by western blotting; qPCR was carried out on samples depleted of protein for 24 or 48 h.

### Quantitative RT-PCR

Total RNA was isolated using Qiagen RNeasy Plus Mini kit or Pure Link RNA mini kit and resuspended in nuclease-free water. Reverse transcription of 250 – 500ng of RNA was carried out using SuperscriptIV with random hexamers following the manufacturer’s instructions. All RT-PCRs were run on the CFX96 Real-Time System using SYBR green Mastermix (Bio-Rad machine, following this program: RT at, 95°C for 2 min, then 40 cycles of 95°C for 30 sec, 55°C for 20 sec, 70°C for 20 sec followed by the plate read step. Each sample was run in 3 technical replicates. Gene expression data was analysed by the delta Ct method, with each gene normalised to housekeeping gene RNA Polymerase II Subunit J (POL2RJ).

### Cellular Fractionation and RNA Sequencing

HeLa cells were mock-depleted or depleted twice of SEC13 & UPF2 as described above. The sample was subsequently split, and total RNA was isolated from depleted cells using PureLink RNA Mini Kit (Life Technologies) according to manufacturer’s instructions. DNA was removed using TURBO DNA-free™ DNase I kit (Invitrogen Ambion; AM1907). Cellular fractionation was performed on the rest of the sample as described previously (Jagannathan et al. 2011), with some modifications. Briefly, four days after the first depletion, cells were detached using trypsin and washed twice in 1ml of ice-cold PBS (500g, 10 min, 4°C). Cellular pellets were resuspended in 0.4 ml of permeabilization buffer (110 mM KOAc, 25 mM K-HEPES pH 7. 2, 2.5 mM Mg(OAc)2, 1 mM EGTA, 0.015% digitonin, 1mM DTT, 1× Complete Protease Inhibitor Cocktail, 40 U/mL RNaseOUT™) and incubated for 5 min on a rotating wheel at 4°C. The resulting cytosolic fractions were recovered by centrifugation at 2,000g for 10 min, 4°C. Cells were then lysed in 0.4 ml of NP-40 lysis buffer (400 mM KOAc, 25 mM K-HEPES pH 7.2, 15 mM Mg(OAc)2, 1% (v/v) NP-40, 1 mM DTT, 1× Complete Protease Inhibitor Cocktail, 40 U/mL RNaseOut) for 30 min on ice. The membrane fraction was recovered by centrifugation at 7,000g for 10 min at 4°C. Both cytosolic and membrane fractions were clarified by centrifugation at 7,500g for 10 min at 4°C. Digitonin, DTT, Complete Protease Inhibitor Cocktail and RNaseOUT™ were added fresh to the buffers. RNA from both fractions were isolated using PureLink RNA Mini Kit (Life Technologies) according to manufacturer’s instructions. DNA was removed using TURBO DNA-free™ DNase I kit (Invitrogen Ambion; AM1907).

### RNA-sequencing analysis

#### Read processing and alignment

Raw RNA-sequencing reads were processed using the nf-core/rnaseq pipeline (v3.21.0) (Ewels et al. 2020) on the University of Edinburgh Eddie HPC cluster. Reads were trimmed with Trim Galore (v0.6.10) (https://www.bioinformatics.babraham.ac.uk/projects/trim_galore/) using Cutadapt (v4.9)((Martin 2011), aligned to the human reference genome (GRCh38, Ensembl release 115) using STAR (v2.7.11b) (Dobin et al. 2013), and quantified at the transcript-level with Salmon (v1.10.3; (Patro et al. 2017). Gene-level counts were derived from Salmon output using tximeta/tximport (v1.20.1) (Soneson et al. 2015), with length scaling to correct for changes in average transcript length across samples. The resulting count matrix and sample metadata were used for all downstream analyses.

#### Quality control

All analyses were conducted in R (v4.3.1 or later) using Bioconductor (v3.18 or later). Gene annotations were retrieved from Bioconductor’s AnnotationHub package using the Ensembl 111 EnsDb for Homo sapiens (record AH119325), linking Ensembl gene IDs to gene symbols, Entrez IDs, and biotype metadata. Initial quality control was performed on the count data. Sample relationships and potential outliers or batch effects were assessed by Principal Component Analysis (PCA) on variance-stabilized expression data generated using the vst function from the DESeq2 package (Love et al. 2014). Potential confounding variables were evaluated by testing correlations between sample metadata and principal components explaining at least 5% of the variance.

#### Differential expression analysis

Differential expression analysis was performed using DESeq2 (v1.46.0). The design formula modeled the main experimental factor of interest, and log2 fold change (LFC) shrinkage was performed using the ashr method to moderate the LFCs of genes with low expression or high dispersion, providing more accurate estimates for ranking and visualization. Genes with adjusted p-value < 0.01 were considered significantly differentially expressed. Differential expression was performed independently for Total RNA, Cytoplasmic RNA, and Membrane-bound RNA. For Total RNA, pairwise contrasts were computed between each knockdown (*SEC13, UPF2, SEC23, SEH1L, WDR24, NUP133*) and the non-targeting control (NTC). For the fractionated samples, pairwise contrasts were computed between each knockdown and NTC within each fraction (Cytoplasmic and Membrane), and between the Membrane and Cytoplasmic fractions within NTC to establish a baseline subcellular localization reference. Results were visualized using several standard plots. Volcano plots, created with the EnhancedVolcano package, were used to visualize the relationship between statistical significance (-log10(padj)) and magnitude of change (LFC). Heatmaps of significant genes, generated with the pheatmap package, were used to display expression patterns across samples. Expression levels of the top differentially expressed genes were visualized using boxplots created with ggplot2 (Wickham 2016).

#### Pathway enrichment analysis and DEG overlap

Reactome pathway over-representation analysis was performed with ReactomePA (Ragueneau et al. 2026) using Entrez gene IDs for genes significantly upregulated in SEC13 KD total RNA (padj < 0.01). The background universe was defined as all genes in the SEC13 DESeq2 results with a non-NA Entrez ID. Pathways with adjusted p-value < 0.05 were considered significant. Overlap among upregulated genes was visualized using an area-proportional Euler diagram. SEC13 and UPF2 knockdown datasets from this study were compared with the UPF1 knockdown dataset from Longman et al. 2020 (GEO: GSE152437). Upregulated genes were defined as adjusted p-value < 0.01 and LFC < 0 in the NTC-vs-KD convention used here, or LFC >0 and adjusted p-value < 0.01 for the published UPF1 dataset.

#### LFC correlation between SEC13 and UPF2 knockdowns

To characterize the correlation of gene changes between SEC13 and UPF2 knockdowns, genes significant at padj < 0.01 in total RNA comparisons were matched by Ensembl gene ID, and per-gene log2 fold changes (KD vs NTC) were plotted against each other. Plots were displayed using the central 95% of values on each axis to improve visualisation. Quadrant counts reflect all both-significant genes including those outside the zoomed window. Pearson correlation coefficients and two-sided p-values were calculated, and the 20 genes with the smallest adjusted p-value in the nominated knockdown were labeled.

#### DESeq2 interaction model for Membrane/Cytoplasm partitioning

Per-gene Membrane vs Cytoplasm (Mem/Cyto) partitioning was quantified using a DESeq2 interaction model fitted to raw counts from the Membrane and Cytoplasmic fraction samples for NTC, SEC13, and UPF2 conditions (∼ fraction + condition + fraction: condition). Genes with fewer than 10 counts in at least 3 samples were excluded. Mem/Cyto log2 fold changes for each condition were estimated from the fitted model coefficients,and the per-gene localisation shift upon SEC13 KD (Δ LFC) was calculated as the difference between the SEC13 and NTC Mem/Cyto LFCs.

#### Antibodies

Western Blotting: Anti-UPF1 (# A300-036/ A300-38A, Bethyl), Anti-GAPDH (#ab9483, Abcam); Anti-Sec13 (# 15397-1-AP, Proteintech); Anti-UPF2 (# sc-374230 Santa Cruz Biotechnology); Anti-UPF3 (# sc-48800 Santa Cruz Biotechnology); Anti-SMG7 (# A302-170A, Bethyl); Anti-Flag (#F3165 Sigma); Anti- βActin (#A5441 Sigma). Imaging: Anti-Sec61β (# 15087-1-AP, Proteintech); Anti-UPF1 (#A300-38A, Bethyl); Anti-Sec13 (# 15397-1-AP, Proteintech). For Immunoprecipitations GFP-Trap-MA beads (Chromotek) were used.

PLA: Anti-GFP (#ab290, Abcam/ 11814460001, Roche); Anti-Sec61β (# 15087-1-AP, Proteintech); Anti-UPF2 (# sc-374230 Santa Cruz Biotechnology); Anti-UPF1 (#A300-38A, Bethyl).

### Statistical analysis

Prism Version 11 software (GraphPad) was used for statistical analysis. Data are displayed as the mean ± s.e.m. Statistical analysis was performed using unpaired two-tailed t-test for comparison of two groups or one-way ANOVA for 3 or more groups. RStudio was used for the statistical analysis of RNA sequencing analysis. For information about the number of replicates, see the corresponding figure legend. For information about how data was analyzed and/or quantified, see the relevant section in Materials and methods.

## Data availability

The following secure token has been created to allow review of record GSE329468 while it remains in private status: admpwomejpmfhqn https://eur02.safelinks.protection.outlook.com/?url=https%3A%2F%2Fwww.ncbi.nlm.nih.gov%2Fgeo%2 Fquery%2Facc.cgi%3Facc%3DGSE329468&data=05%7C02%7C%7Cd0781d61f5574c70093908dea787 62e2%7C2e9f06b016694589878910a06934dc61%7C0%7C0%7C639132397095929727%7CUnknown%7CTWFpbGZsb3d8eyJFbXB0eU1hcGkiOnRydWUsIlYiOiIwLjAuMDAwMCIsIlAiOiJXaW4zMiIsIkFO IjoiTWFpbCIsIldUIjoyfQ%3D%3D%7C0%7C%7C%7C&sdata=c7iT47z%2BBIabvw05KOUyaGTTJGy jUmq9lAJgbN1CTbA%3D&reserved=0

## ACKNOWLEDGMENTS

This work was supported by MRC University Unit grant MC_UU_00035/5. We thank Tom Deegan (MRC HGU) for help with AlphaFold structural modeling. We acknowledge Jimi Wills of the Mass Spectrometry Facility, Matthew Pearson and Laura Murphy of Advanced Imaging resource, Elizabeth Freyer and Michael Rennie of the Flow Cytometry facility, and the Bioinformatics Analysis Core, all at the Institute of Genetics and Cancer (IGC), for their excellent technical support.

## Author contributions

L.M, D.L. and J.F.C. conceived and designed the project and analyzed the data. L.M. performed most of the experimental work, A.M. contributed to the generation of tagged cell lines, and interactome experiments. C.S. and D.L developed the NMD sensors. G.G. carried out the bioinformatic analysis. The manuscript was co-written by all authors.

## SUPPLEMENTAL MATERIAL

Supplemental material is available for this article.

## REFERENCES

Abramson J, Adler J, Dunger J, Evans R, Green T, Pritzel A, Ronneberger O, Willmore L, Ballard AJ, Bambrick J, et al. 2024. Accurate structure prediction of biomolecular interactions with AlphaFold 3. Nature 630: 493–500.

Acosta-Alvear D, Harnoss JM, Walter P, Ashkenazi A. 2025. Homeostasis control in health and disease by the unfolded protein response. Nature reviews Molecular cell biology 26: 193–212.

Anastasaki C, Longman D, Capper A, Patton EE, Cáceres JF. 2011. Dhx34 and Nbas function in the NMD pathway and are required for embryonic development in zebrafish. Nucleic acids research 39: 3686–94.

Applequist SE, Selg M, Raman C, Jäck HM. 1997. Cloning and characterization of HUPF1, a human homolog of the Saccharomyces cerevisiae nonsense mRNA-reducing UPF1 protein. Nucleic acids research 25: 814–21.

Barlowe C, Orci L, Yeung T, Hosobuchi M, Hamamoto S, Salama N, Rexach MF, Ravazzola M, Amherdt M, Schekman R. 1994. COPII: a membrane coat formed by Sec proteins that drive vesicle budding from the endoplasmic reticulum. Cell 77: 895–907.

Bhuvanagiri M, Schlitter AM, Hentze MW, Kulozik AE. 2010. NMD: RNA biology meets human genetic medicine. The Biochemical journal 430: 365–77.

Boehm V, Kueckelmann S, Gerbracht JV, Kallabis S, Britto-Borges T, Altmüller J, Krüger M, Dieterich C, Gehring NH. 2021. SMG5-SMG7 authorize nonsense-mediated mRNA decay by enabling SMG6 endonucleolytic activity. Nature Communications 12: 3965.

Boehm V, Wallmeroth D, Wulf PO, Popp O, Teixeira Alves LG, Reinecke L, Riedel M, Wyler E, Franitza M, Becker K, et al. 2025. Rapid UPF1 depletion illuminates the temporal dynamics of the NMD-regulated human transcriptome. Molecular cell 85: 3524–3546.e12.

Brown WJ, Plutner H, Drecktrah D, Judson BL, Balch WE. 2008. The lysophospholipid acyltransferase antagonist CI-976 inhibits a late step in COPII vesicle budding. Traffic 9: 786–797.

Casadio A, Longman D, Hug N, Delavaine L, Vallejos Baier R, Alonso CR, Cáceres JF. 2015. Identification and characterization of novel factors that act in the nonsense-mediated mRNA decay pathway in nematodes, flies and mammals. EMBO reports 16: 71–8.

Castello A, Hentze MW, Preiss T. 2015. Metabolic Enzymes Enjoying New Partnerships as RNA-Binding Proteins. Trends in endocrinology and metabolism 26: 746–57.

Copley SD. 2014. An evolutionary perspective on protein moonlighting. Biochem Soc Trans 42: 1684–1691.

Dobin A, Davis CA, Schlesinger F, Drenkow J, Zaleski C, Jha S, Batut P, Chaisson M, Gingeras TR. 2013. STAR: Ultrafast universal RNA-seq aligner. Bioinformatics 29: 15–21.

Eden E, Navon R, Steinfeld I, Lipson D, Yakhini Z. 2009. GOrilla: A tool for discovery and visualization of enriched GO terms in ranked gene lists. BMC Bioinformatics 10: 48

Efstathiou S, Ottens F, Schütter LS, Ravanelli S, Charmpilas N, Gutschmidt A, Le Pen J, Gehring NH, Miska EA, Bouças J, et al. 2022. ER-associated RNA silencing promotes ER quality control. Nature cell biology 24: 1714–1725.

Embree CM, Abu-Alhasan R, Singh G. 2022. Features and factors that dictate if terminating ribosomes cause or counteract nonsense-mediated mRNA decay. Journal of Biological Chemistry 298: 102592.

Enninga J, Levay A, Fontoura BMA. 2003. Sec13 shuttles between the nucleus and the cytoplasm and stably interacts with Nup96 at the nuclear pore complex. Molecular and cellular biology 23: 7271–7284.

Ewels PA, Peltzer A, Fillinger S, Patel H, Alneberg J, Wilm A, Garcia MU, Di Tommaso P, Nahnsen S. 2020. The nf-core framework for community-curated bioinformatics pipelines. Nat Biotechnol 38: 276–278.

Fritz SE, Ranganathan S, Wang CD, Hogg JR. 2022. An alternative UPF1 isoform drives conditional remodeling of nonsense-mediated mRNA decay. The EMBO journal 41: e108898.

Gadella TWJ, van Weeren L, Stouthamer J, Hink MA, Wolters AHG, Giepmans BNG, Aumonier S, Dupuy J, Royant A. 2023. mScarlet3: a brilliant and fast-maturing red fluorescent protein. Nature methods 20: 541–545.

Goetz AE, Wilkinson M. 2017. Stress and the nonsense-mediated RNA decay pathway. Cellular and Molecular Life Sciences 74: 3509–3531.

Gopalsamy A, Bennett EM, Shi M, Zhang W-GG, Bard J, Yu K. 2012. Identification of pyrimidine derivatives as hSMG-1 inhibitors. Bioorganic & medicinal chemistry letters 22: 6636–6641.

Gowravaram M, Bonneau F, Kanaan J, Maciej VD, Fiorini F, Raj S, Croquette V, Le Hir H, Chakrabarti S. 2018. A conserved structural element in the RNA helicase UPF1 regulates its catalytic activity in an isoform-specific manner. Nucleic Acids Research 46: 2648–2659.

Hartmann E, Sommer T, Prehn S, Görlich D, Jentsch S, Rapoport TA. 1994. Evolutionary conservation of components of the protein translocation complex. Nature 367: 654–657.

Hollien J, Lin JH, Li H, Stevens N, Walter P, Weissman JS. 2009. Regulated Ire1-dependent decay of messenger RNAs in mammalian cells. The Journal of Cell Biology 186: 323–331.

Huang L, Lou C-H, Chan W, Shum EY, Shao A, Stone E, Karam R, Song H-W, Wilkinson MF. 2011. RNA homeostasis governed by cell type-specific and branched feedback loops acting on NMD. Molecular cell 43: 950–61.

Hug N, Cáceres JF. 2014. The RNA Helicase DHX34 Activates NMD by promoting a transition from the surveillance to the decay-inducing complex. Cell Reports 8: 1845–1856.

Hughes H, Budnik A, Schmidt K, Palmer KJ, Mantell J, Noakes C, Johnson A, Carter DA, Verkade P, Watson P, et al. 2009. Organisation of human ER-exit sites: Requirements for the localisation of Sec16 to transitional ER. Journal of Cell Science 122: 2924–2934.

Iwawaki T, Akai R, Kohno K, Miura M. 2004. A transgenic mouse model for monitoring endoplasmic reticulum stress. Nature Medicine 10: 98–102.

Jagannathan S, Hsu JCC, Reid DW, Chen Q, Thompson WJ, Moseley AM, Nicchitta CV. 2014. Multifunctional roles for the protein translocation machinery in rna anchoring to the endoplasmic reticulum. Journal of Biological Chemistry 289: 25907–25934.

Jagannathan S, Nwosu C, Nicchitta CV. 2011. Analyzing mRNA localization to the endoplasmic reticulum via cell fractionation. *Methods in molecular biology (Clifton*, NJ) 714: 301–21.

Karam R, Lou C-HC-H, Kroeger H, Huang L, Lin JH, Wilkinson MF. 2015. The unfolded protein response is shaped by the NMD pathway. EMBO reports 16: 599–609.

Karousis ED, Mühlemann O. 2022. The broader sense of nonsense. Trends in biochemical sciences 47: 921–935.

Kashima I, Yamashita A, Izumi N, Kataoka N, Morishita R, Hoshino S, Ohno M, Dreyfuss G, Ohno S. 2006. Binding of a novel SMG-1-Upf1-eRF1-eRF3 complex (SURF) to the exon junction complex triggers Upf1 phosphorylation and nonsense-mediated mRNA decay. Genes & development 20: 355–67.

Kim YK, Maquat LE. 2019. UPFront and center in RNA decay: UPF1 in nonsense-mediated mRNA decay and beyond. RNA 25: 407–422.

Kolakada D, Campbell AE, Galvis LB, Li Z, Lore M, Jagannathan S. 2024. A system of reporters for comparative investigation of EJC-independent and EJC-enhanced nonsense-mediated mRNA decay. Nucleic acids research 52:e34.

Kurosaki T, Popp MW, Maquat LE. 2019. Quality and quantity control of gene expression by nonsense-mediated mRNA decay. Nature reviews Molecular cell biology 20: 406–420.

Le Hir H, Gatfield D, Izaurralde E, Moore MJ. 2001. The exon-exon junction complex provides a binding platform for factors involved in mRNA export and nonsense-mediated mRNA decay. The EMBO journal 20: 4987–97.

Loh B, Jonas S, Izaurralde E. 2013. The SMG5-SMG7 heterodimer directly recruits the CCR4-NOT deadenylase complex to mRNAs containing nonsense codons via interaction with POP2. Genes & development 27: 2125–2138.

Longman D, Hug N, Keith M, Anastasaki C, Patton EE, Grimes G, Cáceres JF. 2013. DHX34 and NBAS form part of an autoregulatory NMD circuit that regulates endogenous RNA targets in human cells, zebrafish and Caenorhabditis elegans. Nucleic acids research 41: 8319–31.

Longman D, Jackson-Jones KA, Maslon MM, Murphy LC, Young RS, Stoddart JJ, Hug N, Taylor MS, Papadopoulos DK, Cáceres JF. 2020. Identification of a localized nonsense-mediated decay pathway at the endoplasmic reticulum. Genes & development 34: 1075–1088.

Longman D, Plasterk RHA, Johnstone IL, Cáceres JF. 2007. Mechanistic insights and identification of two novel factors in the C. elegans NMD pathway. Genes & development 21: 1075–85.

Lord C, Ferro-Novick S, Miller EA. 2013. The highly conserved COPII coat complex sorts cargo from the endoplasmic reticulum and targets it to the golgi. Cold Spring Harbor perspectives in biology 5: a013367.

Love MI, Huber W, Anders S. 2014. Moderated estimation of fold change and dispersion for RNA-seq data with DESeq2. Genome Biology 15: 550.

Martin M. 2011. Cutadapt removes adapter sequences from high-throughput sequencing reads. EMBnet.journal 17: 10–12.

McMahon M, Maquat LE. 2025. Exploring the therapeutic potential of modulating nonsense-mediated mRNA decay. RNA 31: 333–348.

Monaghan L, Longman D, Cáceres JF. 2023. Translation-coupled mRNA quality control mechanisms. EMBO J 42: e114378.

Nazarenus T, Cedarberg R, Bell R, Cheatle J, Forch A, Haifley A, Hou A, Kebaara BW, Shields C, Stoysich K, et al. 2005. Upf1p, a highly conserved protein required for nonsense-mediated mRNA decay, interacts with the nuclear pore proteins Nup100p and Nup116p. Gene 345: 199–212.

Ottens F, Efstathiou S, Hoppe T. 2024. Cutting through the stress: RNA decay pathways at the endoplasmic reticulum. Trends in Cell Biology 34: 1056–1068.

Paillusson A, Hirschi N, Vallan C, Azzalin CM, Mühlemann O. 2005. A GFP-based reporter system to monitor nonsense-mediated mRNA decay. Nucleic Acids Research 33: e54.

Papandreou I, Denko NC, Olson M, Van Melckebeke H, Lust S, Tam A, Solow-Cordero DE, Bouley DM, Offner F, Niwa M, et al. 2011. Identification of an Ire1alpha endonuclease specific inhibitor with cytotoxic activity against human multiple myeloma. Blood 117: 1311–1314.

Patro R, Duggal G, Love MI, Irizarry RA, Kingsford C. 2017. Salmon provides fast and bias-aware quantification of transcript expression. Nature Methods 14: 417–419.

Pereverzev AP, Gurskaya NG, Ermakova GV, Kudryavtseva EI, Markina NM, Kotlobay AA, Lukyanov SA, Zaraisky AG, Lukyanov KA. 2015. Method for quantitative analysis of nonsense-mediated mRNA decay at the single cell level. Scientific reports 5: 7729.

Ragueneau E, Gong C, Sinquin P, Sevilla C, Beavers D, Grentner A, Griss J, Hogue GFJ, Li NT, Matthews L, et al. 2026. The Reactome Knowledgebase 2026. Nucleic Acids Res 54: D673–D681.

Ran FA, Hsu PD, Wright J, Agarwala V, Scott DA, Zhang F. 2013. Genome engineering using the CRISPR-Cas9 system. Nature protocols 8: 2281–2308.

Reid DW, Nicchitta CV. 2015. Diversity and selectivity in mRNA translation on the endoplasmic reticulum. Nature Reviews Molecular Cell Biology 16: 221–231.

Schindelin J, Arganda-Carreras I, Frise E, Kaynig V, Longair M, Pietzsch T, Preibisch S, Rueden C, Saalfeld S, Schmid B, et al. 2012. Fiji: An open-source platform for biological-image analysis. Nature Methods 9: 676–682.

Serin G, Gersappe A, Black JD, Aronoff R, Maquat LE. 2001. Identification and characterization of human orthologues to Saccharomyces cerevisiae Upf2 protein and Upf3 protein (Caenorhabditis elegans SMG-4). Molecular and cellular biology 21: 209–23.

Shaywitz DA, Orci L, Ravazzola M, Swaroop A, Kaiser CA. 1995. Human SEC13Rp functions in yeast and is located on transport vesicles budding from the endoplasmic reticulum. The Journal of cell biology 128: 769–77.

Sieber J, Hauer C, Bhuvanagiri M, Leicht S, Krijgsveld J, Neu-Yilik G, Hentze MW, Kulozik AE. 2016. Proteomic analysis reveals branch-specific regulation of the unfolded protein response by nonsense-mediated mRNA decay. Molecular & cellular proteomics 15: 1584–97.

Siniossoglou S, Wimmer C, Rieger M, Doye V, Tekotte H, Weise C, Emig S, Segref A, Hurt EC. 1996. A novel complex of nucleoporins, which includes Sec13p and a Sec13p homolog, is essential for normal nuclear pores. Cell 84: 265–275.

Soneson C, Love MI, Robinson MD. 2015. Differential analyses for RNA-seq: transcript-level estimates improve gene-level inferences. F1000Res 4: 1521.

Stagg SM, Gürkan C, Fowler DM, LaPointe P, Foss TR, Potter CS, Carragher B, Balch WE. 2006. Structure of the Sec13/31 COPII coat cage. Nature 439: 234–8.

Tan K, Wilkinson MF. 2025. Biological roles of nonsense-mediated RNA decay: insights from the nervous system. Current Opinion in Genetics and Development 93: 102356.

Trcek T, Sato H, Singer RH, Maquat LE. 2013. Temporal and spatial characterization of nonsense-mediated mRNA decay. Genes & development 27: 541–51.

Valenstein ML, Rogala KB, Lalgudi PV, Brignole EJ, Gu X, Saxton RA, Chantranupong L, Kolibius J, Quast JP, Sabatini DM. 2022. Structure of the nutrient-sensing hub GATOR2. Nature 607: 610–616.

Walter P, Ron D. 2011. The unfolded protein response: from stress pathway to homeostatic regulation. Science 334: 1081–1086.

Wegener M, Dietz KJ. 2022. The mutual interaction of glycolytic enzymes and RNA in post-transcriptional regulation. RNA 28: 4446–4468.

Weng Y, Czaplinski K, Peltz SW. 1996. Genetic and biochemical characterization of mutations in the ATPase and helicase regions of the Upf1 protein. Molecular and cellular biology 16: 5491–506.

Whittle JRR, Schwartz TU. 2010. Structure of the Sec13-Sec16 edge element, a template for assembly of the COPII vesicle coat. Journal of Cell Biology 190: 347–361.

Wickham H. 2016. *ggplot2: Elegant Graphics for Data Analysis*. Springer-Verlag New York https://ggplot2.tidyverse.org.

Yepiskoposyan H, Aeschimann F, Nilsson D, Okoniewski M, Mühlemann O. 2011. Autoregulation of the nonsense-mediated mRNA decay pathway in human cells. RNA 17: 2108–18.

